# Mitochondrial function determines severity but not risk of amyotrophic lateral sclerosis

**DOI:** 10.1101/2022.05.31.494229

**Authors:** Calum Harvey, Marcel Weinreich, Sai Zhang, Paul J Hop, Ramona A J Zwamborn, Kristel van Eijk, Thomas H Julian, Tobias Moll, Alfredo Iacoangeli, Ahmad Al Khleifat, John P Quinn, Abigail L Pfaff, Sulev Koks, Joanna Poulton, Stephanie L Battle, Dan E Arking, Michael P Snyder, Project MinE ALS Sequencing Consortium, Jan Veldink, Kevin P Kenna, Pamela J Shaw, Johnathan Cooper-Knock

## Abstract

Amyotrophic lateral sclerosis (ALS) is a fatal neurodegenerative disease. Selective vulnerability of energy-intensive motor neurons (MNs) has fostered speculation that mitochondrial function is a determinant of ALS. Previously, the position of mitochondrial function in the pathogenic cascade leading to neurotoxicity has been unclear. We separated upstream genetic determinants of mitochondrial function, including genetic variation within the mitochondrial genome or autosomes; from downstream changeable factors including mitochondrial copy number (mtCN) and MN gene expression. We discovered that functionally validated mitochondrial haplotypes are a determinant of ALS survival but not ALS risk. Loss-of-function genetic variants within, and reduced MN expression of, *ACADM* and *DNA2* lead to shorter ALS survival; both genes impact mitochondrial function. MtCN responds dynamically to the onset of ALS independent of mitochondrial haplotype, and is also significantly correlated with disease severity. We conclude that mitochondrial function impacts ALS progression but not risk; our findings have therapeutic implications.

## Background

Amyotrophic lateral sclerosis (ALS) is a rapidly progressive, relatively common and incurable neurodegenerative disease. In the majority of cases, ALS is thought to result from a complex interaction of genetic and environmental risk factors^1^. The development of ALS is distinct from its progression. Whilst disease can take >50 years to develop, paralysis as a result of motor neuron (MN) loss can occur within <2 years from symptom onset^2^. It is interesting that the genetic basis of ALS risk and ALS survival are largely distinct^3^. Indeed, specific genetic^4^ and environmental factors^5^ have been associated with severity of the clinical phenotype.

Inadequate mitochondrial energy production has long been associated with neurodegenerative disease^6–8^, but questions have remained regarding the position of energy deficit in the pathophysiological cascade^9^. It has been difficult to establish whether energy deficit is an upstream determinant of disease, a downstream consequence of neurodegeneration, or an adjunct which accelerates neuronal death caused by other factors. Primary mitochondrial disease is associated with sensorimotor neuropathy, but specific toxicity to motor neurons is not a cardinal feature^10^. However, several lines of evidence exist for specific links between ALS pathology and mitochondrial dysfunction: TDP-43 mislocalization, which is the hallmark of >97% of ALS^11^, is associated with TDP-43 entry into the mitochondria and impaired bioenergetic function^12^. Moreover, in limbic-predominant age-related TDP-43 encephalopathy (LATE), TDP-43 pathology has been associated with reduced mitochondrial DNA copy number (mtCN) in cortical tissue^13^. Treatments for ALS aimed at reducing the bioenergetic deficit have so far proved ineffective^14^.

One difficulty in studying mitochondrial energy production is the method employed for measurement of bioenergetic function. MtCN is a measure of the number of mitochondria within a biological sample. Higher copy number corresponds to a greater number of mitochondria and potentially a greater capacity for energy production^15^, although this relationship is not linear^16^. However, MtCN varies throughout life and adapts to the particular metabolic needs of each specific cell and cell-type^17^. MtCN can be modified by environmental factors including the onset of disease and ageing^18^, which makes it difficult to differentiate the effect of mtCN on disease from the effect of disease on mtCN. In contrast, mitochondrial DNA sequence is determined largely by maternal inheritance which is fixed at conception and as a result mitochondrial haplotypes are necessarily upstream of a late age of onset disease such as ALS. We took advantage of mitochondrial haplotypes which have been robustly associated with mtCN, with mtCN-associated quantitative traits, and with non-cancer mortality^19^. Our objective was to use mitochondrial haplotypes as a measure of upstream mitochondrial function independent of the effect of disease onset. We tested for an effect of mitochondrial haplotype on ALS risk and/or survival. Multiple number of nuclear encoded genes have also been associated with mitochondrial function^20^ and genetic variation within nuclear encoded genes is also largely fixed at conception; therefore, we performed a gene burden analysis as an orthogonal test of a causal link between mitochondrial function and ALS risk and/or survival.

Our data are not consistent with an upstream role for mitochondrial function as a determinant of the risk of developing ALS. However, we do find evidence for modification of ALS survival by genetically-determined mitochondrial function, suggesting that this may be a valid therapeutic target for a majority of ALS patients. Our findings have important implications for efforts to understand the role of energy deficit in the pathophysiology of ALS. Our approach is summarised in **Figure 1**.

**Figure 1:**
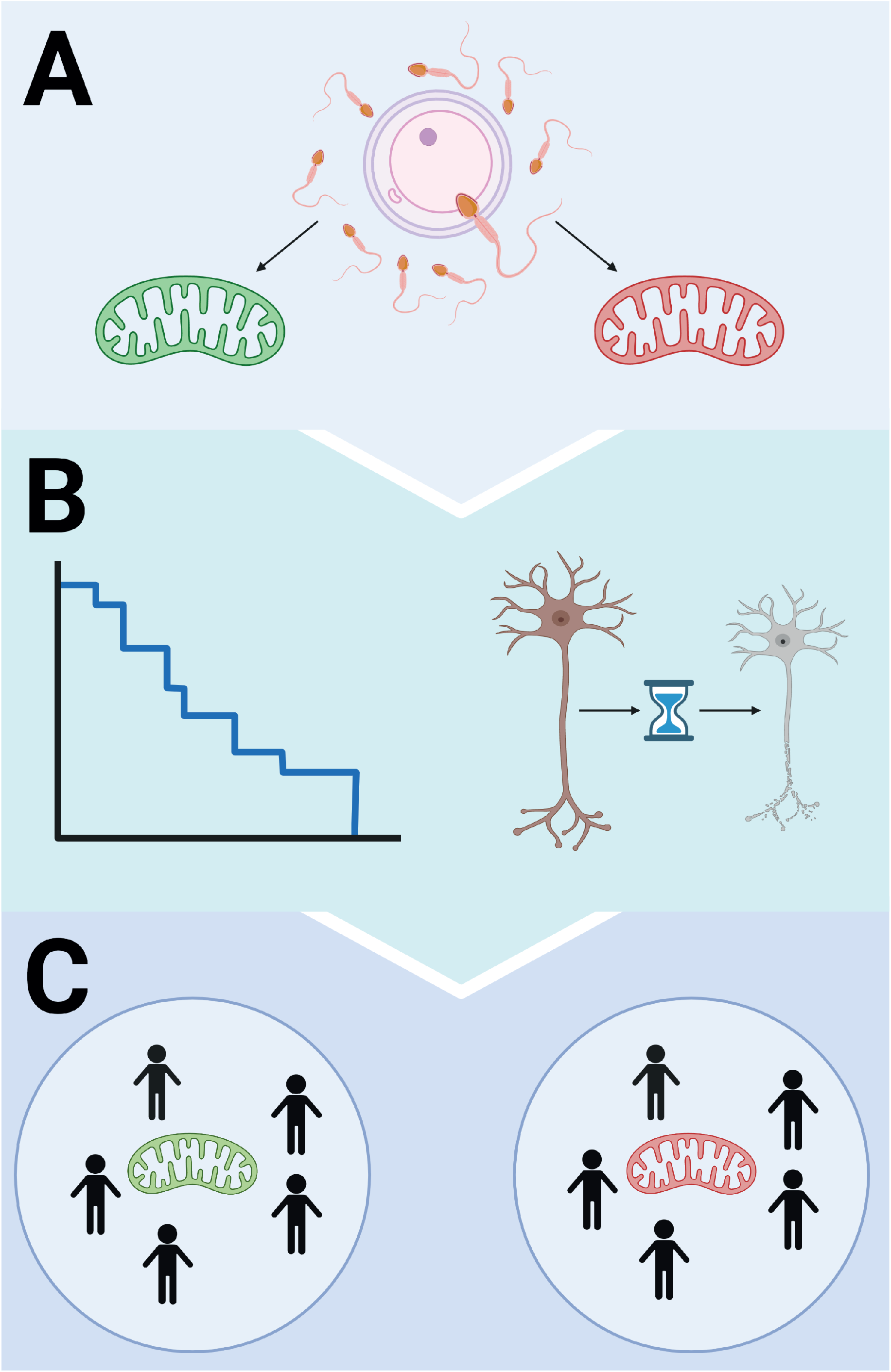
Genetic determinants of mitochondrial function are used to test for a causal role of bioenergetic function in ALS risk and progression. Schematic of the study design. (a) Mitochondrial haplotypes and genetic variants impacting mitochondrial function are fixed at conception. (b) ALS development, progression and survival are downstream of genetic determinants of mitochondrial function. (c) Comparison between groups with differing mitochondrial haplotypes and/or genetic variants can therefore be made to understand how mitochondrial function affects ALS development, progression and survival.

## Methods

### Study cohort

The 5,594 sporadic unrelated ALS patients and 2,238 controls subject to WGS and included in this study as part of the Project MinE cohort were recruited at specialized neuromuscular centres in the UK, Belgium, Germany, Ireland, Italy, Spain, Turkey, the United States and the Netherlands^21^. Patients were diagnosed with possible, probable or definite ALS according to the 1994 El-Escorial criteria^22^. All controls were free of neuromuscular diseases and matched for age, sex and geographical location. This included sequencing of DNA obtained from whole blood, but also lymphoblastoid cells derived from 896 ALS patients and 400 controls recruited in the UK. Measurement of the ALS functional rating scale (ALSFRS)^23^ was available for at least one timepoint in 1,576 patients.

The AnswerALS cohort (https://www.answerals.org/) consisted of WGS in 843 ALS patients and 15 controls recruited from specialised neuromuscular centres in the United States. For a subset of 180 of these ALS patients, we obtained transcriptome data from iPSC-derived MN, and ALSFRS measurements.

The study was approved by the South Sheffield Research Ethics Committee. Also, this study followed study protocols approved by Medical Ethical Committees for each of the participating institutions. Written informed consent was obtained from all participating individuals. All methods were performed in accordance with relevant national and international guidelines and regulations.

### Sample quality control

The Project MinE WGS dataset is recruited from a large number of different countries and healthcare settings. ALS survival is affected by access to healthcare and specific care delivered^24^. Kaplan-Meier curve analysis revealed heterogeneity in survival between Project MinE cohorts (**Supplementary Figure 1a**). Based on this analysis Turkish and Portuguese cohorts were excluded from further analysis.

### Mitochondrial haplotype

Mitochondrial haplogroups considered were not exhaustive, but were assigned based on six mitochondrial SNPs selected because they have significant, independent associations with mtCN associated traits^19^. All ALS and control samples were assigned to a particular haplotype. One haplotype was present in <2.5% of ALS samples and therefore this haplotype was excluded from further analysis. The relative frequencies of all haplotypes are shown in **Supplementary Figure 1b**. In subsequent analyses the most common haplogroup was set as the reference and significance for haplogroup associations was generated by comparison to the reference group.

The six mitochondrial SNPs used are MT73A_G, MT7028C_T, MT10238T_C, MT7028C_T, MT12612A_G, MT13617T_C, MT15257G_A (revised Cambridge reference sequence, reference allele first, alternate second). Haplogroups are displayed as ordered MT73_MT7028_MT10238_MT12612_MT13617_MT15257, with 2 as the reference allele, 0 as the alternate allele e.g. 2_2_2_2_2_2 would be the reference allele at every position whereas 0_2_2_2_2_2 would have the alternate allele for SNP MT73.

### Mitochondrial DNA copy number

In recent times whole genome sequencing (WGS) has been established as the gold standard method for measuring mtCN^25^ due to high depth (>2000X) coverage over the mitochondrial genome. MtCN was estimated using the ratio of reads aligning to mitochondrial DNA: the mtCN was calculated as double the number of reads aligning to a set of non-repetitive regions throughout the autosome. These non-repetitive regions were identified as control regions for the profiling of repeat expansions^26^ and were transposed to build 37 for use with the Project MINE data using the UCSC liftover tool.

### White blood cell proportions

White blood cell (WBC) proportions including granulocytes, monocytes, natural killer (NK) cells, T-cells (CD4+ and CD8+) and B cells were estimated from methylation data in the Project MinE cohort as previously described^27^. Platelets counts were not available as they cannot be inferred from methylation data which may impact on the estimation of mtCN because platelets contain mitochondrial DNA but no autosomal DNA^16^.

### Transcriptome analysis

For AnswerALS data, gene expression profiling of iPSC-derived MNs and phenotype data were obtained for 180 ALS patients (https://www.answerals.org/). Gene expression was normalized for gene length and then sequencing depth to produce transcripts per kilobase million (TPM). We used multivariable linear regression to determine the relationship between gene expression and rate of change in the ALSFRS using platform, sex, site of onset (spinal or bulbar), *C9ORF72* status, and the first 20 principal components of genetic variation were included as covariates. Significance testing was performed for all genes (n=30,807) expressed in MNs as determined by mean TPM>1. In this unbiased analysis no gene was significantly associated after multiple testing correction but there was no evidence of inflation or deflation in the test statistics (**Supplementary Figure 2**).

### Statistical analysis

All time to event analyses including survival and age of ALS onset were conducted using a Cox proportional hazards model. Analysis of the relationship between mitochondrial haplotype and mtCN utilised multivariable linear regression. Analysis of the relationship between MN gene expression or mtCN and ALSFRS-slope utilised multivariable linear regression. Analysis of the relationship between mitochondrial haplotype or mtCN and ALS status was conducted by multivariable logistic regression. All analyses included age, sex, site of disease onset, the first 20 principal components of genetic variation, and sequencing platform as co-variates. Additional co-variates are specified in the relevant sections. Sequencing platform was not available for the AnswerALS dataset.

We measured seven mitochondrial haplogroups which were considered as categorical variables; each individual haplotype was compared to the most frequent haplogroup 2_2_2_2_2_2, which was designated as the reference. We report the results of these analyses in our multivariable comparisons but to assess the significance of haplotype overall we performed an ANOVA between regression models with and without the consideration of mitochondrial haplotypes. ANOVA enables comparison of the deviance between the model fits based on the log partial likelihood.

In survival analysis, left truncation bias^28^ was corrected by considering only the time between patient sampling and time of death. Left truncation bias occurs when risk of death is measured over a time in which it could not have occurred; by definition a patient who had already died could not have been recruited into a study. The rationale is to exclude survival time that occurred before sampling, because the patient could not, by definition, have died during this period. Analysis of left truncated data can lead to false positive associations^28^, however, left truncation bias correction can lead to under-powered analysis because of the loss of information. As a result, we report both results for analysis in both the Project MinE and AnswerALS cohorts.

### Rare variant association testing

For analysis of WGS data from 5,594 sporadic ALS patients and 2,238 controls^21^, variants within coding regions were determined to be rare if the minor allele frequency (MAF) within the Genome Aggregation Database (gnomAD) is <1/100 control alleles^29^. In coding regions, we annotated variants using Variant Effect Predictor (VEP)^30^; Loss-of-function (LoF) variants were defined as nonsense mutations, high-effect splice-site mutations^31^, or 5’UTR variants involving a gain/loss of a start/stop codon^32^. We applied a Cox proportional hazards model to determine the relationship between the number of rare LoF variants per patient and disease survival or censored survival time. Sequencing platform, sex, site of onset (spinal or bulbar), *C9ORF72* status, and the first 20 principal components of genetic variation were included as covariates. To test for a link between the number of rare LoF variants per patient and risk of ALS we used Firth logistic regression as previously described^33^. We excluded tests including <5 variants as these are likely to be underpowered.

## Results

### Mitochondrial genotype is an upstream determinant of ALS survival but not ALS risk

We took advantage of mitochondrial haplotypes which have been robustly associated with mtCN, with mtCN-associated quantitative traits, and with disease^19^ (**Methods**). We infer that these mitochondrial haplotypes are associated with mitochondrial function. By dividing ALS and control genomes based on mitochondrial haplotype we were able to explore whether mitochondrial function is associated with both ALS risk and/or survival. We used WGS of DNA extracted from 5,635 sporadic ALS patients and 2,270 controls^21^ (**Methods**).

We first set out to validate the link between mitochondrial haplotypes and mtCN in our dataset. MtCN is itself a complex phenotype determined by interaction of host genetics and the environment^34^. MtCN is cell-specific^18^ and ALS is known to alter the proportions of blood leukocytes^3^. To overcome this limitation we used a subset of our whole blood WGS cohort for whom we could derive proportions of WBC based on DNA methylation^27^ (**Methods**); this subset included 3,549 ALS patients and 1,529 controls. As expected, specific mitochondrial haplotypes were associated with mtCN in our dataset after adjusting for WBC proportions (0_0_2_2_2_2 p=0.007, beta=6.12, se=2.26; 0_2_2_2_2_2, p=0.05, beta=9.01, se=4.65, multivariable linear regression, **Figure 2a, Supplementary Table 1, Methods**); our findings match a previous study of the same haplotypes in peripheral blood^19^.

**Figure 2:**
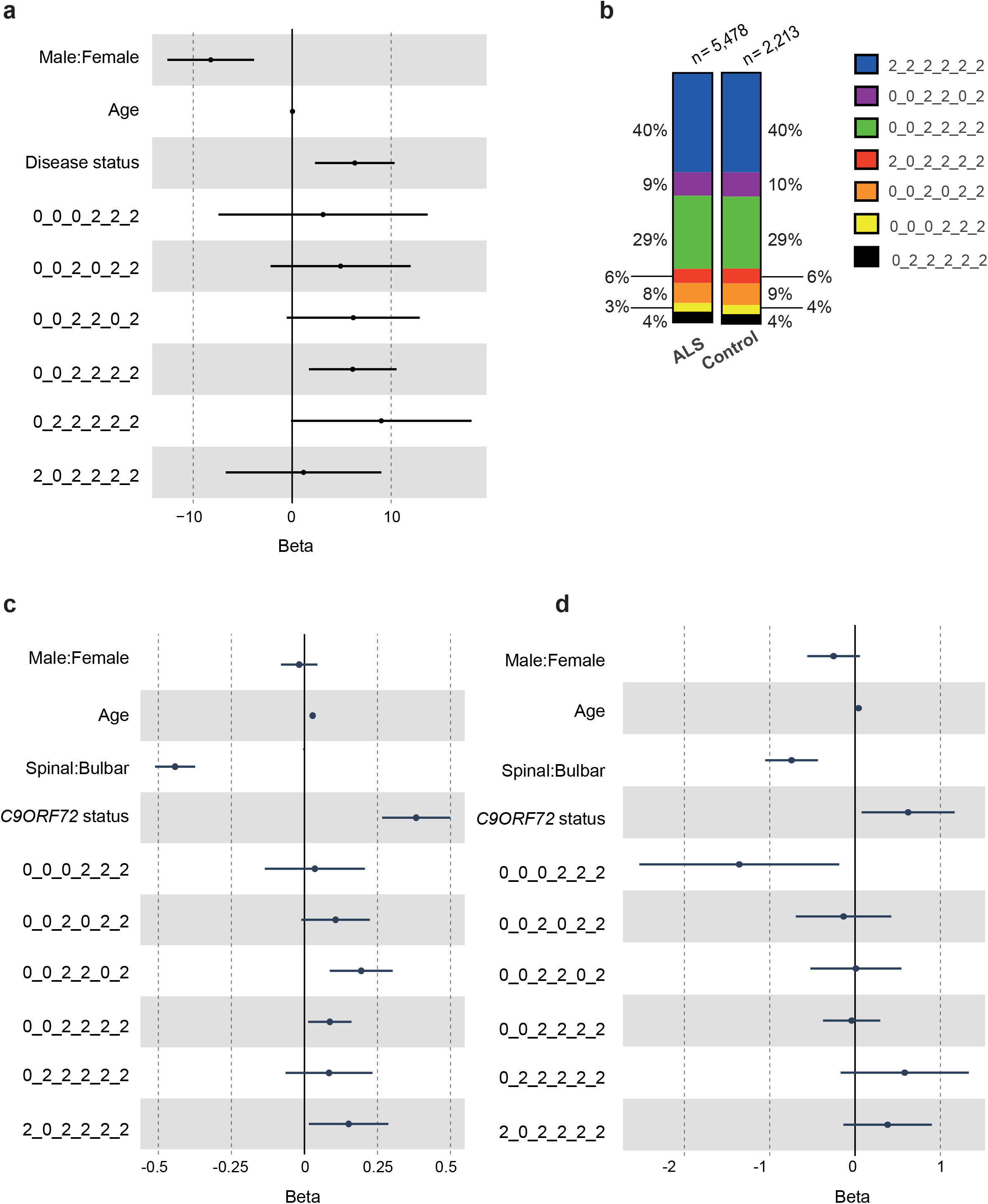
Mitochondrial haplotype is linked to mitochondrial copy number and ALS survival but not ALS risk. (a) Forest plot for the effect of mitochondrial haplotype, sex, age and disease status on mitochondrial copy number. (b) Relative proportions of measured mitochondrial haplotypes amongst ALS patients and controls. (c) Forest plot for the effect of mitochondrial haplotype, sex, age, site of onset and *C9ORF72* status on ALS survival in samples from the Project MinE cohort (n=5,635). (d) Forest plot for the effect of mitochondrial haplotype, sex, age, site of onset and *C9ORF72* status on ALS survival in samples from the AnswerALS cohort (n=843). Mitochondrial haplotypes were determined by six mitochondrial SNPs: MT73A_G, MT7028C_T, MT10238T_C, MT7028C_T, MT12612A_G, MT13617T_C, MT15257G_A. Haplogroups are displayed as ordered MT73_MT7028_MT10238_MT12612_MT13617_MT15257, with 2 as the reference allele, 0 as the alternate allele.

We did not find any evidence that mitochondrial haplotype was associated with ALS risk (p>0.05, logistic regression, **Figure 2b, Methods**). Surprisingly, mitochondrial haplotype was significantly associated with ALS survival (p=0.0096, Chisq=16.91, ANOVA). Three individual mitochondrial haplogroups were significantly associated with ALS survival (0_0_2_2_0_2, p=4.1e-4, HR=1.21, 95%CI: 1.09-1.35; 0_0_2_2_2_2, p=0.022, HR=1.09, 95%CI: 1.01-1.18; and 2_0_2_2_2_2, p=0.029, HR=1.16 95%CI: 1.02-1.33; Cox regression, **Figure 2c, Supplementary Table 2, Methods**) and 0_0_2_2_0_2 was significant after stringent Bonferroni multiple testing correction. After correction of left truncation bias (**Methods**) mitochondrial haplotype is still significantly associated with ALS survival (p=0.023, Chisq=14.7, ANOVA) and the deleterious effect of 0_0_2_2_0_2 (p=7.3e-3, HR=1.19, 95%CI: 1.05-1.34) and 0_0_2_2_2_2 (p=0.01, HR=1.12, 95%CI: 1.03-1.22) is maintained; 0_0_2_2_0_2 remained significant after Bonferroni multiple testing correction (**Supplementary Figure 3a, Supplementary Table 3, Methods**). In our analysis we excluded diagnostic delay (time from symptom onset to diagnosis) as a covariate because of some concern that this may be determined in part by mitochondrial haplotype; interestingly when diagnostic delay is included as a covariate then the effect of mitochondrial haplotype is actually enhanced (p=1.9e-04, Chisq=26.3, ANOVA, **Supplementary Figure 3b, Supplementary Table 4, Methods**) suggesting that mitochondrial haplotype has an outsized effect on survival time after diagnosis. Survival data included 4,549 patients who had died and 930 censored data points where the patient is still alive.

Mitochondrial haplotype is fixed at conception and is therefore necessarily upstream of a late age of onset disease such as ALS. Our data suggests that mitochondrial dysfunction is not a risk factor for ALS but mitochondrial function can affect the progression of ALS once disease has been initiated.

### Mitochondrial genotype is associated with ALS survival in an independent cohort

To validate our findings we obtained WGS data from an additional cohort including 843 ALS patients (https://www.answerals.org/, **Methods**). This cohort was enriched with long survivors compared to the test cohort (median survival 3.15 years versus 2.72 years). In this analysis mitochondrial haplotype was still significantly associated with ALS survival (p=0.047, Chisq=12.7, ANOVA). Consistent with the longer survival in this group the most protective mitochondrial haplotype reached statistical significance in isolation (0_0_0_2_2_2 p=0.023, HR=0.26, 95%CI: 0.08-0.83; Cox regression, **Figure 2d, Supplementary Table 5, Methods**) and this remained significant after correction for left truncation bias (0_0_0_2_2_2 p=0.026, HR=0.26, 95%CI: 0.08-0.852, **Supplementary Figure 3c, Supplementary Table 6, Methods**). Indeed, this protective haplotype was overrepresented in this cohort compared to the original test cohort (**Supplementary Figure 1b**). Direction of effect and effect size for all mitochondrial haplotypes are similar in both cohorts (**Figure 2c-d**).

### Mitochondrial copy number is elevated in ALS patient tissues

We have suggested that mitochondrial function is not an upstream cause of ALS. We next asked whether the initiation of the ALS disease process impacts upon mtCN. We tested for a significant relationship between ALS diagnosis and mtCN after adjusting for WBC proportions and mitochondrial haplotype. Consistent with a change in mtCN in response to disease, mtCN is significantly elevated in ALS patients compared to controls (p=2.03e-3, beta=1.94e-3, se=6.30e-4, multivariable logistic regression), but the effect size is very small (OR=1.004, **Figure 3a, Supplementary Table 7, Methods**).

**Figure 3:**
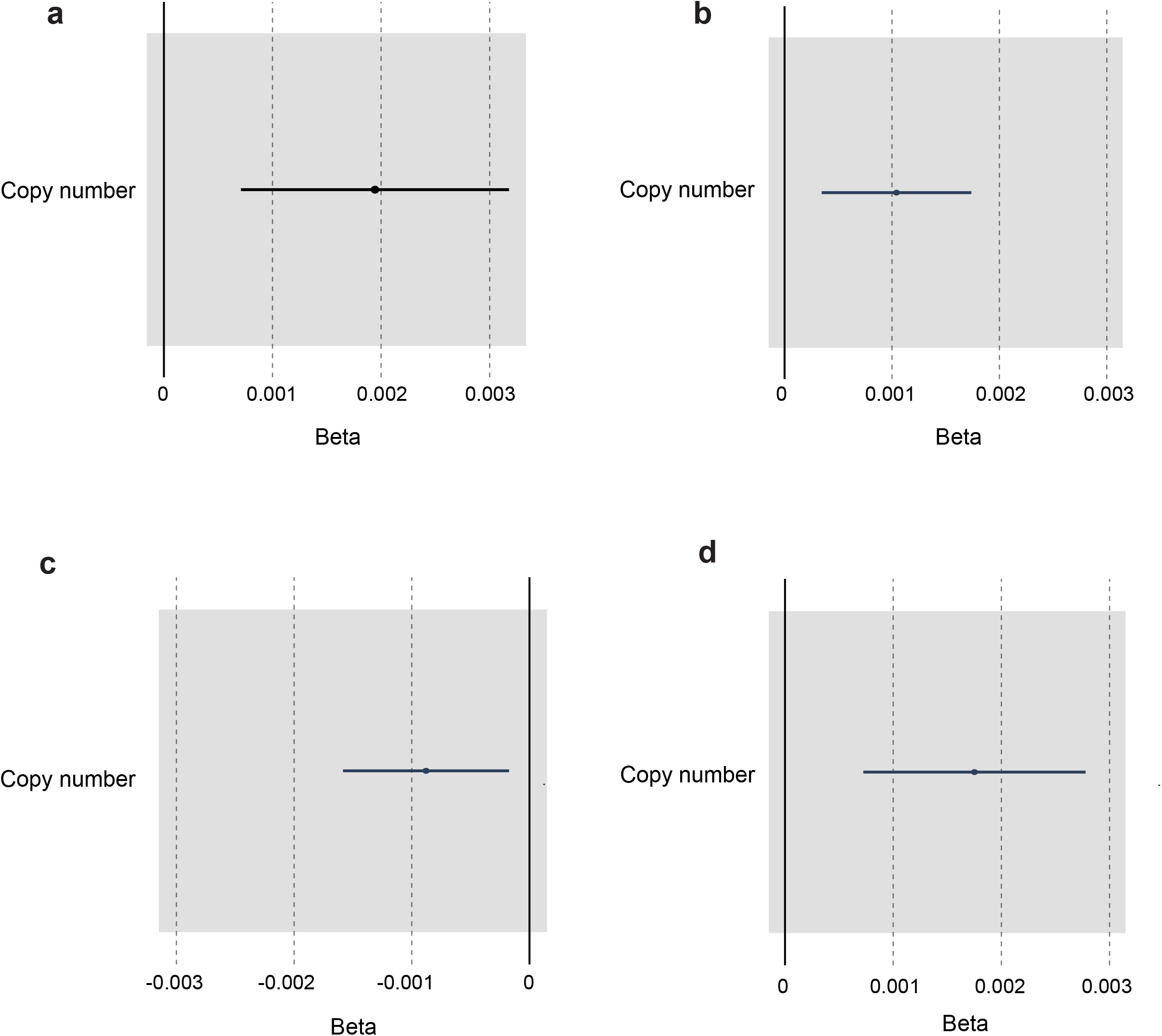
Mitochondrial copy number is linked to ALS survival and age of disease onset. (a) Forest plot for the correlation between mitochondrial copy number and ALS. (b) Forest plot for the correlation between mitochondrial copy number and ALS survival. (c) Forest plot for the correlation between mitochondrial copy number and ALS age of onset. (d) Forest plot for the correlation between mitochondrial copy number and age at sampling in controls.

Taken together with our analysis of mitochondrial haplotype, this suggests that changes in mtCN may occur downstream of disease initiation in an attempted compensation for neuronal stress or for mislocalization of TDP-43 to the mitochondria^12^.

### Mitochondrial copy number is associated with ALS severity

We have shown that mitochondrial function is an upstream modifier of ALS severity. Moreover, our analysis of mtCN suggests that the onset of ALS itself may precipitate an increase in mtCN. We hypothesised that changes in mtCN may be proportional to disease severity independent of the effect of mitochondrial haplotype. For example patients who achieve a higher mtCN may be more likely to avoid neurotoxic bioenergetic stress, or else a more aggressive disease course may precipitate a greater increase in mtCN. To test this hypothesis we examined whether mtCN was correlated with disease severity within our WGS dataset after correcting for both WBC proportions and mitochondrial haplotype. In this analysis higher mtCN is significantly associated with shorter ALS survival (p=0.0037, HR=1.001, 95%CI: 1-1.002, Cox regression, **Figure 3b, Supplementary Table 8, Methods**).

### The link between mitochondrial copy number and age is modified by ALS

Age is negatively correlated with mtCN measured in peripheral blood^19^. If ALS is a consistent modifier of mtCN throughout the disease course, we questioned whether the normal relationship between mtCN and age might be altered.

Higher mtCN is associated with higher age at blood sampling for ALS patients (p=0.015, HR=0.999, 95%CI 0.998-1, **Figure 3c, Supplementary Table 10**). In contrast, in our control cohort the direction of association is reversed (p=8.3e-04, HR=1.002, 95%CI:1.001-1.003, Cox regression, **Fig 3d, Supplementary Table 11**). This is consistent with previous literature demonstrating lower mtCN with advanced age in normal individuals^19^. We conclude that the biological mechanism mediating changes in mtCN in response to ALS is dynamic over time and is independent of the effects of normal aging. To explore this further we tested for a relationship between mtCN and rate of change in the ALS functional rating scale (ALSFRS)^23^ (**Methods**), which is a measure of ALS progression. However, after adjusting for age there was no evidence of a significant relationship (p=0.61, beta=0.0454, se=0.09, **Supplementary Table 12**).

Mitochondrial haplotype is not significantly associated with ALS age of onset (p=0.24, Chisq=7.9, ANOVA, **Supplementary Figure 3d, Supplementary Table 9, Methods**).

### Function of nuclear encoded genes linked to mitochondrial function impact on ALS survival

Next we analysed MN expression of nuclear encoded genes associated with mitochondrial function to determine whether they might be linked to ALS rate of progression. We obtained a published and curated list of 166 nuclear encoded genes associated with mitochondrial function and used to study neurodegenerative disease^20^. We utilised RNAseq from iPSC-derived motor neurons obtained from 180 ALS patients (https://www.answerals.org/). RNAseq data and survival data were available for only 82 individuals because many of the patients were not followed-up until death. Therefore we focused on the link between motor neuron gene expression and rate of change in the ALSFRS which was available for all 180 patients. Rate of change in the ALSFRS is frequently used to measure ALS progression^23^. We tested for an association between MN expression of nuclear encoded mitochondrial genes with rate of change in the ALSFRS by multivariable linear regression (**Figure 4a, Supplementary Table 13, Methods**) but no gene was significant after Bonferroni multiple testing correction. However, compared to the background transcriptome (**Methods**) the p-value for mitochondrial genes was significantly lower than expected (p=4.4e-05, rank sum test). Moreover, for all genes which achieved nominal significance (p<0.05) expression was inversely related to rate of change in the ALSFRS (**Figure 4a**) meaning that lower expression is linked to increased rate of progression. This was significant compared to the direction of correlation for the background transcriptome (p=5e-4, Fisher Exact test). We conclude from this analysis that mitochondrial function within motor neurons is associated with the rate of ALS progression; this orthogonal validation is consistent with our earlier analysis of mitochondrial haplotype and mtCN.

**Figure 4:**
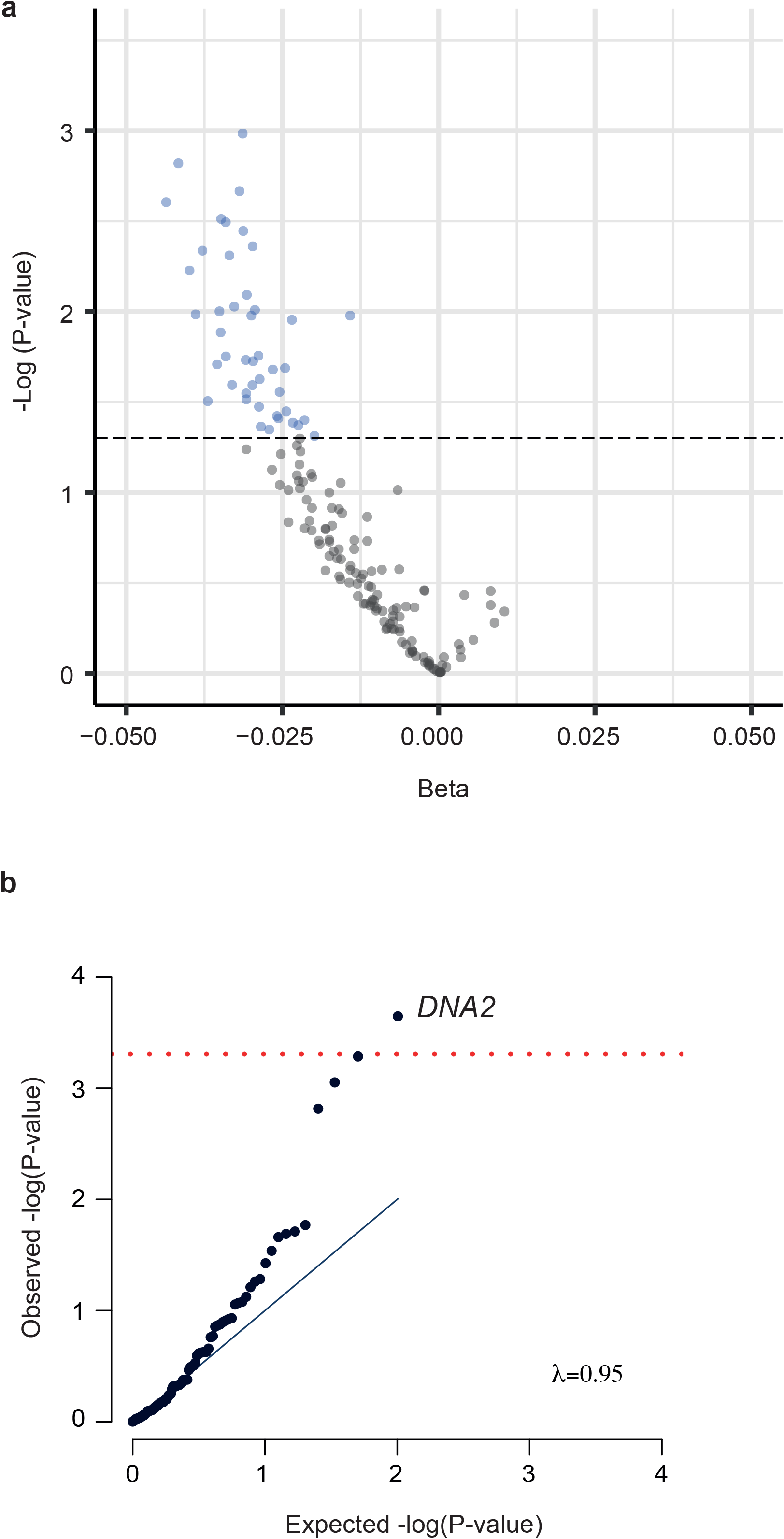
Motor neuron expression of nuclear encoded genes associated with mitochondrial function is associated with rate of disease progression. (a) Volcano plot including effect sizes and test statistics for multivariable linear regression including the effect of expression of nuclear encoded genes associated with mitochondrial function on rate of ALS progression. Black dotted line and blue circles indicate p<0.05. (b) QQ-plot for expected and observed P-values in Cox regression including the effect of burden of rare loss of function genetic variants within nuclear encoded genes associated with mitochondrial function on ALS survival. The test statistics show no evidence of inflation (λ=1.005). One gene, DNA2, is significant after Bonferroni multiple testing (indicated by red line). 166 genes were tested.

Returning to the Project MinE dataset we tested whether LoF genetic variants (**Methods**) within the 166 nuclear encoded genes associated with mitochondrial function^20^, could influence ALS survival. Like mtCN, the motor neuron transcriptome is changeable and potentially downstream of the onset of ALS. However genetic variation within nuclear encoded genes is fixed at conception and therefore necessarily upstream. One gene, *DNA2*, was significantly enriched with LoF mutations in ALS patients with shorter survival after Bonferroni multiple testing correction (p=2.3e-4, beta=1.17, Cox regression, **Figure 4b, Supplementary Table 14, Methods**); mean survival in twelve ALS patients carrying a LoF mutation in *DNA2* was 1.83 years compared to 3.46 years in patients without a LoF mutation within *DNA2*. To leverage statistical power, we performed a meta-analysis of our MN expression and LoF studies which revealed that two genes were significantly associated with ALS severity after Bonferroni multiple testing correction (*ACADM*: p=1.76e-05; and *DNA2*: p=4.44e-05; Stouffer’s test, **Methods**). Both genes are still significant after Bonferroni multiple testing correction even if a stringent correction for left truncation bias is applied (*ACADM*: p=1.46e-04; and *DNA2*: p=2.40e-04; Stouffer’s test, **Methods**). It is notable that for both *ACADM* and *DNA2*, LoF mutations and lower MN gene expression are associated with faster disease progression; moreover both genes are nominally significant in both analyses (**Supplementary Table 13** and **Supplementary Table 14**). Mean survival in four ALS patients carrying a LoF mutation in *ACADM* was 1.39 years compared to 3.46 years in patients without a LoF mutation within *ACADM*. LoF mutations within *DNA2* or *ACADM* are not associated with risk for ALS (p>0.5, Firth logistic regression, **Methods**).

## Discussion

ALS is a result of specific toxicity to motor neurons which leads to death, usually from respiratory failure, within 2-5 years^2^. The majority of ALS cases are thought to be the product of a gene-environment interaction^1^. In all cases described to date, genetic risk is present from conception but disease does not develop until much later; most patients do not develop symptoms until their 6th decade. The contrast between the decades it takes to develop disease and the relatively few years it takes for disease to spread through the CNS^2^ has led to the suggestion that these two processes are biologically distinct. In this study we have treated disease risk and disease progression as separate entities and have sought to examine the relationship of each to mitochondrial dysfunction (**Figure 1**).

The role of bioenergetics in neurodegenerative disease is controversial and has been the subject of a number of failed clinical trial interventions for ALS, e.g. the phase 3 dexpramipexole trial^14^. Significant evidence exists from both *in vitro*^*35*^ and *in vivo*^*36*^ studies suggesting that insufficient energy production exacerbates or even causes neurotoxicity. Mitochondria are the source of cellular ATP production and, down-regulation of mitochondrial function at disease onset is associated with a more severe phenotype in the *SOD1*-ALS mouse model^37^. What has been missing until now is a study of pre-morbid mitochondrial function across a large number of ALS patients to determine the effect on disease risk and severity.

To isolate the upstream role of mitochondrial function we used mitochondrial genetic haplotypes with a validated link to mtCN, mitochondria-determined phenotypes and to disease^19^. Our approach is analogous to Mendelian randomization (MR) but traditional MR for mtCN is prohibited by high instrument pleiotropy and the limited power of mtCN-associated common SNPs^19^. Mitochondrial haplotypes are fixed at conception and therefore our analysis is not impacted by secondary effects of disease.

We did not observe a significant link between mitochondrial haplotype and risk of ALS which suggests that mitochondrial function is not a primary cause of the selective degeneration of motor neurons seen in ALS. This is useful information for the field and suggests that therapeutic intervention to modify mitochondrial function is not appropriate in, for example, asymptomatic family members of ALS patients who harbour disease-causing mutations. We have identified a significant relationship between mitochondrial haplotype and ALS survival which we validated in an independent cohort. Our analyses suggest that interventions to modify mitochondrial function may be able to slow disease progression and increase survival time. We provide evidence that mtCN in blood changes dynamically through the course of ALS independently of mitochondrial haplotype. MtCN was elevated in the blood of ALS patients and significantly correlated with survival but interpretation is difficult, particularly because mtCN is tissue specific^17^ and ALS is a disease of the CNS.

The relationship between mitochondrial haplotype and ALS survival was stronger when we corrected for diagnostic delay, which is the time from symptom onset to diagnosis. Shorter diagnostic delay is typically associated with more rapid disease progression^38^. Our results suggest that this information is non-overlapping with the information provided by mitochondrial haplotype and therefore we infer that mitochondrial haplotype selectively impacts survival time later in the disease course. Our understanding of ALS disease progression as a combination of multiple independent and interacting processes is developing, but isolating the influence of mitochondrial function may delineate specific biological pathways and processes^39^.

Our analysis of mitochondrial haplotype and mtCN has established an upstream role for mitochondrial function in ALS disease progression but the design of therapeutic interventions will require more specific biological insights. We aimed to achieve this through study of patient-derived MNs. Our transcriptome analysis in iPSC-derived MN from ALS patients revealed that the levels of expression of nuclear encoded genes associated with mitochondrial function are significantly associated with the rate of ALS progression. By combining this analysis with rare variant association testing using WGS we were able to associate reduced ALS survival time with LoF mutations within two genes. First *ACA*DM, which encodes the medium-chain specific (C4 to C12 straight chain) acyl-Coenzyme A dehydrogenase. Increased expression of this protein has been identified in the spinal cord of G93A-SOD1-ALS mice^40^. This enzyme is key in the process of fatty acid beta-oxidation for energy production. Importantly, medium-chain fatty acids are able to bypass the carnitine shuttle which is necessary for import of long chain fatty acids into mitochondria; defects in the carnitine shuttle have previously been associated with ALS^41^. A genetic defect leading to deficient metabolism of medium-chain fatty acids might exacerbate this defect by removing a potential escape mechanism. The second mitochondrial gene we identified was *DNA2* which encodes a DNA Replication Helicase/Nuclease 2 which is involved in DNA repair, particularly within the mitochondria. Biallelic variants within *DNA2* have been associated with a developmental phenotype^42^ but our data suggest that heterozygous LoF changes in this gene may impair the bioenergetic response to ALS leading to more rapid disease progression, perhaps via excessive DNA damage within the mitochondrial genome.

A limitation of our results is that our estimate of mtCN failed to include platelet counts because this information was not available. We note that despite this, we were able to replicate previous results including the association of mtCN with specific mitochondrial haplotypes and with age in control individuals^19^.

Overall our work links bioenergetic function within MN to rate of progression of ALS. Equally we have shown that upstream determinants of bioenergetic function are not linked to the risk of developing ALS. Positioning of mitochondrial function as a disease modifier and not as a cause of ALS is vitally important in the search for new therapeutics. It is already known that high caloric diets can improve outcomes in ALS^43^. Targeted intervention aimed at boosting bioenergetic function is likely to be equally or even more effective and potentially synergistic.

## Acknowledgements

This work was supported by the National Institutes of Health (CEGS 5P50HG00773504, 1P50HL083800, 1R01HL101388, 1R01-HL122939, S10OD025212, P30DK116074, and UM1HG009442 to M.P.S.), the Wellcome Trust (216596/Z/19/Z to J.C.-K.), and NIHR (NF-SI-0617-10077 to P.J.S.). This project has received funding from the European Research Council (ERC) under the European Union’s Horizon 2020 research and innovation programme (grant agreement n° 772376 - EScORIAL). AAK is funded by a ALS Association Milton Safenowitz Research Fellowship, The Motor Neurone Disease Association (MNDA) Fellowship and The NIHR Maudsley Biomedical Research Centre. The collaboration project is co-funded by the PPP Allowance made available by Health∼Holland, Top Sector Life Sciences & Health, to stimulate public-private partnerships. This study was also supported by the ALS Foundation Netherlands, by research grants from IWT (n° 140935), the ALS Liga België, the National Lottery of Belgium, the KU Leuven Opening the Future Fund. We also acknowledge support from a Kingsland fellowship (T.M.), and the NIHR Sheffield Biomedical Research Centre for Translational Neuroscience (IS-BRC-1215-20017) and the NIHR Clinical Research Facility. We are very grateful to the ALS patients and control subjects who generously donated biosamples. We acknowledge transcriptomic data provided by the AnswerALS Consortium. Figure 1 was created with BioRender.com.

## Author Contribitions

CH and JCK conceived and designed the study. CH, PJH, RAJZ, SB, DA, JV, PJS and JCK were responsible for data acquisition. CH, MW, THJ, TM, AI, AAK, JQ, ALP, SK, KPK, SZ, PJS and JCK were responsible for analysis of data. CH, MW, THJ, TM, PJH, RAJZ, AI, AAK, JQ, AS, SK, KPK, SZ, JP, SB, DA, MPS, JV, PJS and JCK were responsible for interpretation of data. The Project MinE ALS Sequencing Consortium was involved in data acquisition and analysis. CH, MW, PJS, DEA and JCK prepared the manuscript with assistance from all authors. All authors meet the four ICMJE authorship criteria, and were responsible for revising the manuscript, approving the final version for publication, and for accuracy and integrity of the work.

## Declaration of Interests

M.P.S is a co-founder and member of the scientific advisory board of Personalis, Qbio, January, SensOmics, Protos, Mirvie, NiMo, Onza and Oralome. He is also on the scientific advisory board of Danaher, Genapsys and Jupiter.

## Figure Legends

**Supplementary Figure 1:**
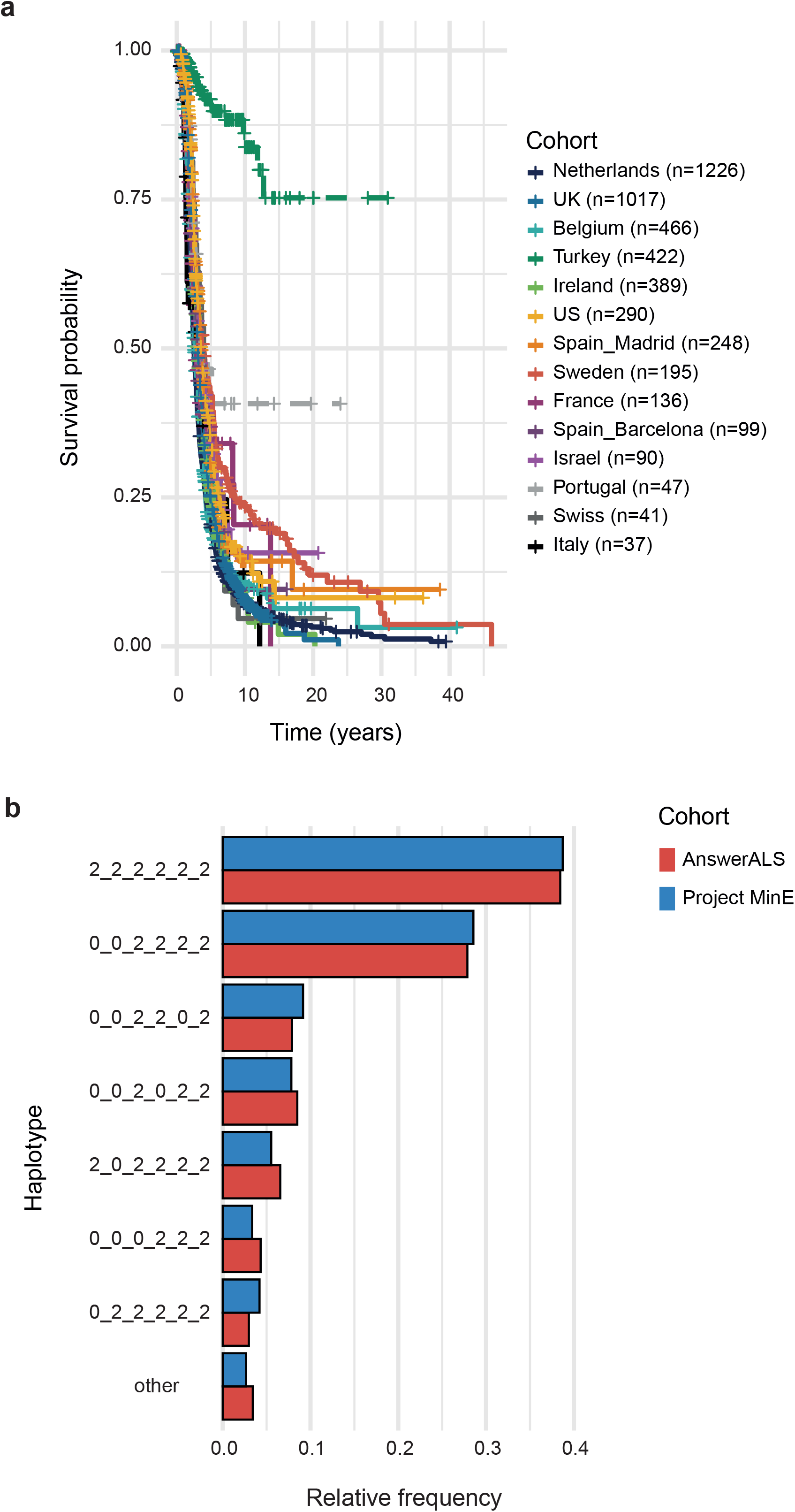
**Survival characteristics and frequency of mitochondrial haplotypes in the Project MinE cohort** including 5,635 sporadic ALS patients. (a) Kaplan-Meier curve revealed significant heterogeneity between subcohorts. As a result of atypical characteristics the Turkish and Portuguese cohorts were excluded from further analysis. Censored survival times are represented by a cross. (b) Relative frequency for each of the studied mitochondrial haplotypes in both the Project MinE and AnswerALS cohorts of ALS patients. 2_2_2_2_2_2 was the most frequent and was taken as the reference haplotype. Only the six haplotypes present in >2.5% of the study cohort were included in further analyses to avoid underpowered tests. Mitochondrial haplotypes were determined by six mitochondrial SNPs: MT73A_G, MT7028C_T, MT10238T_C, MT7028C_T, MT12612A_G, MT13617T_C, MT15257G_A. Haplogroups are displayed as ordered MT73_MT7028_MT10238_MT12612_MT13617_MT15257, with 2 as the reference allele, 0 as the alternate allele.

**Supplementary Figure 2:**
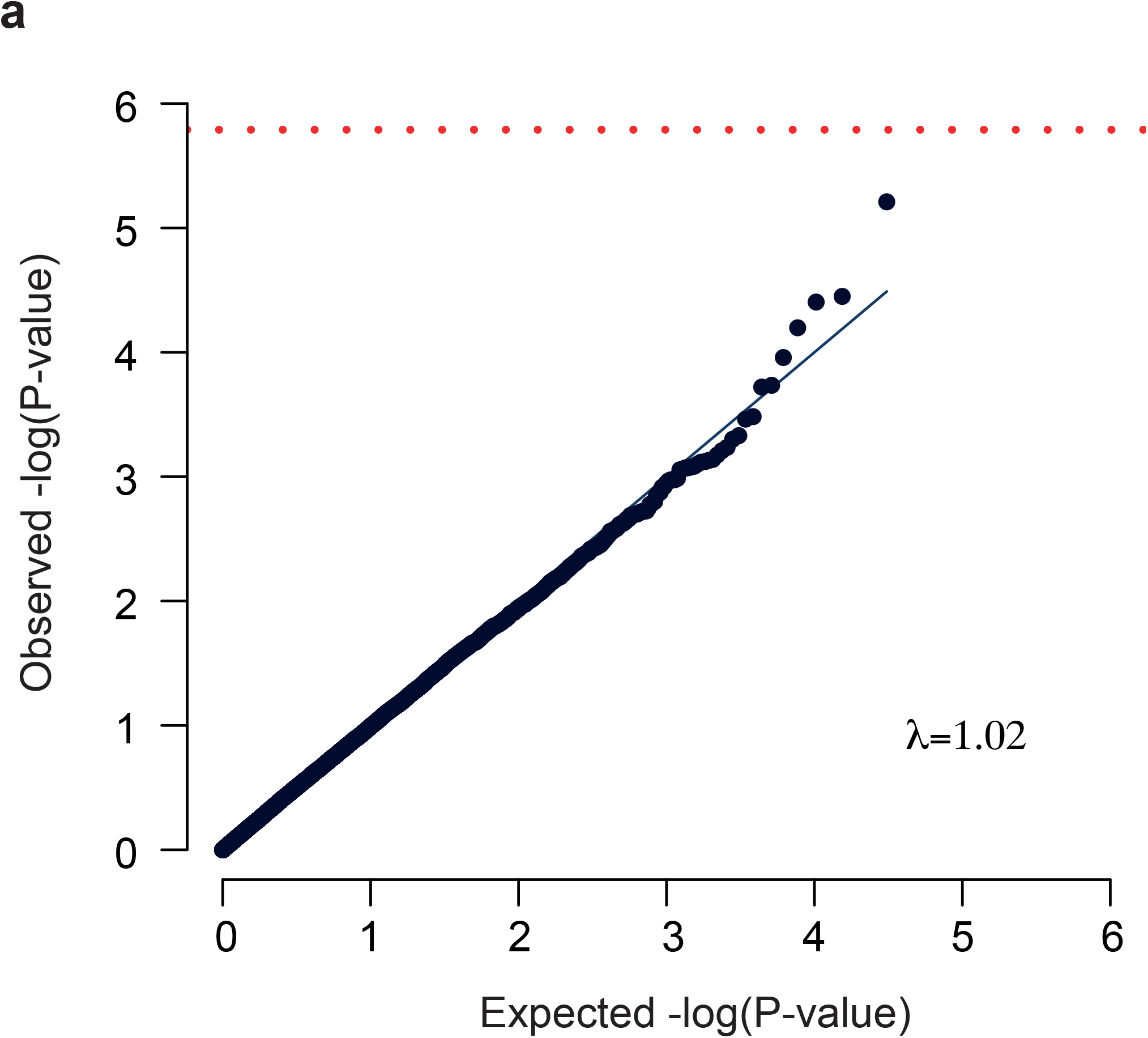
QQ-plot for expected and observed P-values in multivariable linear regression analysis of the effect of gene expression on rate of ALS progression. The test statistics show no evidence of inflation (λ=1.02) but no individual gene is significant after Bonferroni multiple testing (indicated by red line). 30,807 genes were tested. ALS progression was measure by the rate of change in the ALS functional rating score (ALSFRS).

**Supplementary Figure 3:**
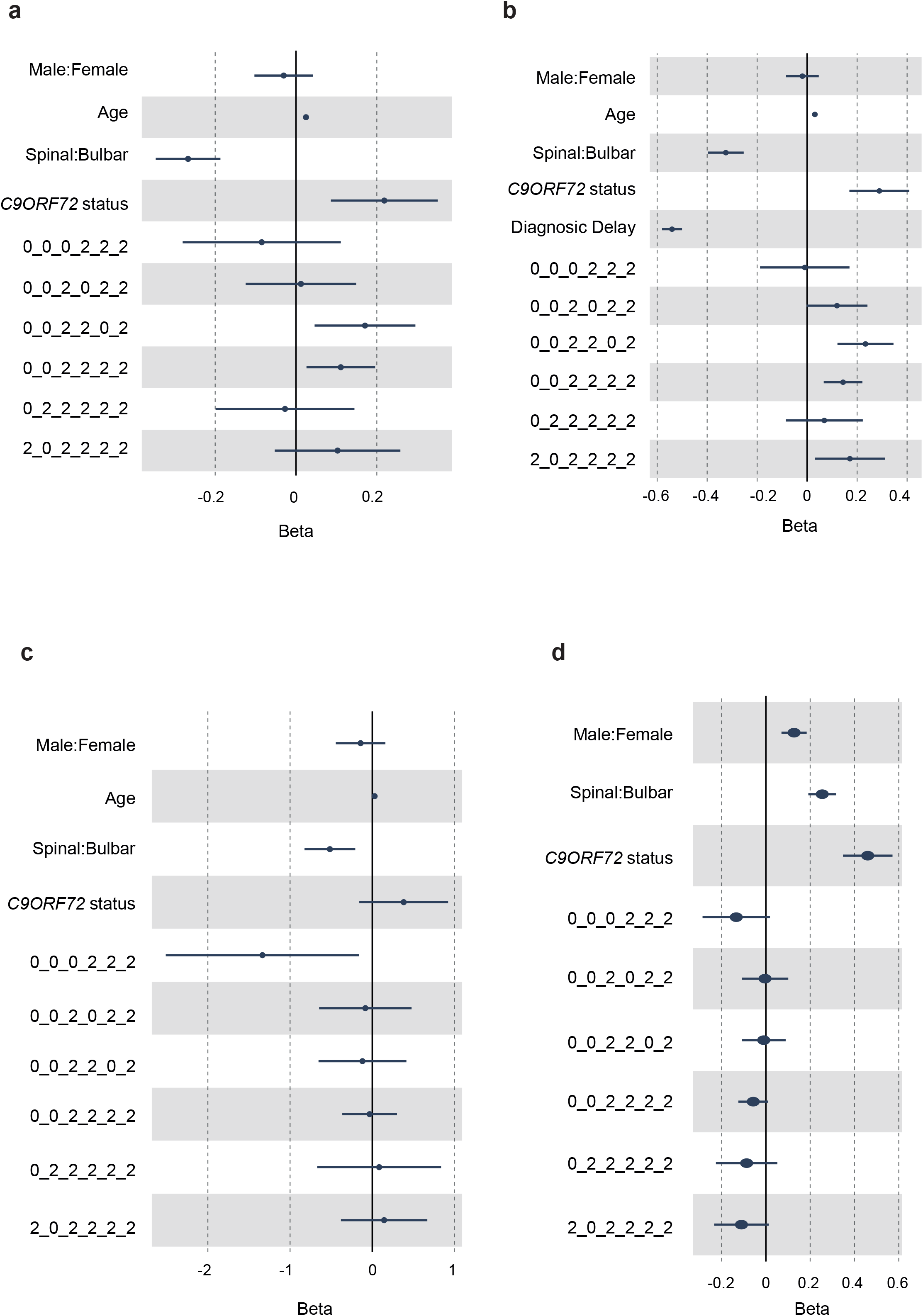
(a) Forest plot for Cox regression penalised to correct for left truncation bias, for the effect of mitochondrial haplotype, sex, age, site of onset and disease status on ALS survival in samples from the Project MinE cohort. (b) Forest plot for the effect of mitochondrial haplotype, sex, age, site of onset, *C9ORF72* status, and diagnostic delay on ALS survival in samples from the Project MinE cohort. (c) Forest plot for Cox regression penalised to correct for left truncation bias, for the effect of mitochondrial haplotype, sex, age, site of onset and disease status on ALS survival in samples from the AnswerALS cohort. (d) Forest plot for the effect of mitochondrial haplotype, sex, site of onset, *C9ORF72* status, on ALS age of onset in samples from the Project MinE cohort.

**Supplementary Table 1: Effect sizes and test statistics for multivariable linear regression including the effect of mitochondrial haplotype on mitochondrial copy number**. This analysis was carried out in blood samples from the Project MinE cohort including 3,549 sporadic ALS patients and 1,529 controls.

**Supplementary Table 2: Effect sizes and test statistics for Cox regression including the effect of mitochondrial haplotype on ALS survival**. This analysis was carried out in samples from the Project MinE cohort including 5,635 sporadic ALS patients.

**Supplementary Table 3: Effect sizes and test statistics for Cox regression penalised to correct for left truncation bias including the effect of mitochondrial haplotype on ALS survival**. This analysis was carried out in samples from the Project MinE cohort including 5,635 sporadic ALS patients.

**Supplementary Table 4: Effect sizes and test statistics for Cox regression including the effect of mitochondrial haplotype and diagnostic delay on ALS survival**. This analysis was carried out in samples from the Project MinE cohort including 5,635 sporadic ALS patients.

**Supplementary Table 5: Effect sizes and test statistics for Cox regression including the effect of mitochondrial haplotype on ALS survival**. This analysis was carried out in the AnswerALS cohort including 843 ALS patients.

**Supplementary Table 6: Effect sizes and test statistics for Cox regression penalised to correct for left truncation bias including the effect of mitochondrial haplotype on ALS survival**. This analysis was carried out in the AnswerALS cohort including 843 ALS patients.

**Supplementary Table 7: Effect sizes and test statistics for multivariable logistic regression including the correlation between mitochondrial copy number and ALS**. This analysis was carried out in blood samples from the Project MinE cohort including 3,549 sporadic ALS patients and 1,529 controls.

**Supplementary Table 8: Effect sizes and test statistics for Cox regression including the correlation between mitochondrial copy number and ALS survival**. This analysis was carried out in blood samples from the Project MinE cohort including 3,549 sporadic ALS patients.

**Supplementary Table 9: Effect sizes and test statistics for Cox regression including the effect of mitochondrial haplotype on ALS age of onset**. This analysis was carried out in samples from the Project MinE cohort including 5,635 sporadic ALS patients.

**Supplementary Table 10: Effect sizes and test statistics for Cox regression including the correlation between mitochondrial copy number and ALS age of onset**. This analysis was carried out in blood samples from the Project MinE cohort including 3,549 sporadic ALS patients.

**Supplementary Table 11: Effect sizes and test statistics for Cox regression including the correlation between mitochondrial copy number and age at sampling in controls**. This analysis was carried out in blood samples from the Project MinE cohort including 1,529 controls.

**Supplementary Table 12: Effect sizes and test statistics for multivariable linear regression including the effect of mitochondrial copy number on rate of ALS progression**. This analysis was carried out in blood samples from the Project MinE cohort including 3,549 sporadic ALS patients.

**Supplementary Table 13: Effect sizes and test statistics for multivariable linear regression including the effect of expression of nuclear encoded genes associated with mitochondrial function on rate of ALS progression**. This analysis was carried out in the AnswerALS cohort including 180 ALS patients.

**Supplementary Table 14: Effect sizes and test statistics for Cox regression including the effect of burden of rare loss of function genetic variants within nuclear encoded genes associated with mitochondrial function on ALS survival**. This analysis was carried out in samples from the Project MinE cohort including 5,635 sporadic ALS patients.

**Supplementary Table 15: Project MinE ALS Sequencing Consortium**

